# Massive influence of DNA isolation and library preparation approaches on palaeogenomic sequencing data

**DOI:** 10.1101/075911

**Authors:** Axel Barlow, Gloria G. Fortes, Love Dalén, Ron Pinhasi, Boris Gasparyan, Gernot Rabeder, Christine Frischauf, Johanna L.A. Paijmans, Michael Hofreiter

## Abstract

The ability to access genomic information from ancient samples has provided many important biological insights. Generating such palaeogenomic data requires specialised methodologies, and a variety of procedures for all stages of sample preparation have been proposed. However, the specific effects and biases introduced by alternative laboratory procedures is insufficiently understood. Here, we investigate the effects of three DNA isolation and two library preparation protocols on palaeogenomic data obtained from four Pleistocene subfossil bones. We find that alternative methodologies can significantly and substantially affect total DNA yield, the mean length and length distribution of recovered fragments, nucleotide composition, and the total amount of usable data generated. Furthermore, we also detect significant interaction effects between these stages of sample preparation on many of these factors. Effects and biases introduced in the laboratory can be sufficient to confound estimates of DNA degradation, sample authenticity and genomic GC content, and likely also estimates of genetic diversity and population structure. Future palaeogenomic studies need to carefully consider the effects of laboratory procedures during both experimental design and data analysis, particularly when studies involve multiple datasets generated using a mixture of methodologies.

## INTRODUCTION

The field of palaeogenomics, defined as the analysis of whole-genomic data from ancient or degraded samples, has emerged relatively recently but has already led to major advances in our understanding of organismal evolution, population genetics, demographics, domestication and human history (reviewed in Shapiro & Hofreiter 2014). The ability to access palaeogenomic data has been largely facilitated by the application of high-throughput sequencing methods, in particular the Illumina sequencing platform, either by direct shotgun sequencing (e.g. Meyer et al. 2012) or by utilising hybridisation capture enrichment for complete mitochondrial (e.g. Paijmans et al. 2015) or nuclear genomes (e.g. Enk et al. 2014). However, the transition from PCR-amplification of individual target sequences to high-throughput sequencing has required significant adaptation and optimisation to the special requirements of palaeogenomic research. For example, it has been shown that standard library construction methods developed for modern DNA perform poorly for ancient DNA, due to the small amounts of starting template generally found in ancient samples, as well as their distinct chemical modifications, leading to the development of specific methods to facilitate the conversion of ancient template molecules into sequencing libraries (Maricic & Pääbo 2009; Meyer *et al.* 2012; Gansauge & Meyer 2013). Similarly, the possibility of high-throughput technology to access extremely short DNA fragments has resulted in modified DNA isolation methods that retain such short molecules (Dabney *et al.* 2013). However, rigorous comparisons of alternative protocols for palaeogenomic research have been relatively infrequent, and studies have generally investigated only one specific stage of sample processing, such as DNA extraction (Rohland & Hofreiter 2007; Gamba *et al.* 2016), library preparation (Dabney & Meyer 2012; Bennett *et al.* 2014; Wales *et al.* 2015) or hybridisation capture (Paijmans *et al.* 2015; Cruz-Dávalos *et al.* 2016; Mohandesan *et al.* 2016). The potential for interactive effects of specific combinations of protocol therefore remains, to our knowledge, almost completely unexplored.

Those studies that have compared alternative protocols for ancient DNA isolation or library preparation have generally focussed on identifying methods that maximise either DNA recovery or the efficiency of data generation (e.g. Rohland & Hofreiter 2007; Bennett et al. 2014; Wales *et al.* 2015; Gamba et al. 2016). The specific effects or biases that particular protocols may introduce to palaeogenomic data are, however, arguably of greater importance because such factors may confound data analysis or interpretation. Parameters of specific interest for palaeogenomic researchers include nucleotide composition and the distribution of DNA fragment lengths. The latter has been used to establish both data authenticity (e.g. Dabney et al. 2013; Hofreiter et al. 2014) and compare levels of degradation among samples (Allentoft et al. 2012). The frequencies of ancient DNA fragments longer than the mode typically decay exponentially. This rate of decay has been modelled mathematically by the lambda parameter, which is assumed to be related to sample age and the thermal deposition environment, and has further been used to estimate the half-life of DNA (Allentoft *et al.* 2012). However, since alternative methods of DNA isolation (Gamba *et al.* 2016) and library preparation (Bennett *et al.* 2014; Wales *et al.* 2015) are known to alter the length distribution of recovered fragments, it stands to reason that laboratory methods will also influence DNA degradation parameters such as lambda, although this has never been directly measured.

In this study, we evaluate the effects of three DNA isolation and two library preparation protocols, as well as their interactive effects, on the quantity and quality of palaeogenomic data recovered from subfossil specimens. We find that these laboratory methods can have substantial and significant effects on total DNA yield, the average length and length distribution of recovered fragments, nucleotide composition, and the total amounts of usable data generated. The implications of these results for palaeogenomic studies are discussed.

## MATERIALS AND METHODS

### Experimental design and laboratory procedures

We compared three DNA isolation and two library preparation protocols using petrous bones from three Late Pleistocene cave bears (*Ursus kudarensis*; samples HV72, HV74 and HV75) and a Late Pleistocene brown bear (*U. arctos*; sample Uap). Full details of samples are provided in Table 1.

**Table 1.** Details of subfossil samples used in this study

| Sample | Taxon | Common name | Age | Locality | Endo %<sup>4</sup> |
| --- | --- | --- | --- | --- | --- |
| HV72 | *U. kudarensis* | Cave bear | 54,000 ($\pm 5700$) yr<sup>1</sup> | Hovk, Armenia | 33.8 |
| HV74 | *U. kudarensis* | Cave bear | 54,000 ($\pm 5700$) yr<sup>1,2</sup> | Hovk, Armenia | 58.4 |
| HV75 | *U. kudarensis* | Cave bear | Late Pleistocene | Hovk, Armenia | 29.5 |
| Uap | *U. arctos* | Brown bear | <sup>14</sup>C cal. age 41,201yr ($\pm 895$yr)<sup>3</sup> | Winden, Austria | 58.7 |

<sup>1</sup>Dated based on stratigraphy (Pinhasi *et al.* 2011)
<sup>2</sup>Sample is beyond range of <sup>14</sup>C dating (>49,000yr). Dating code: MAMS 23142.
<sup>3</sup>Fortes *et al.* 2016.
<sup>4</sup>Endogenous percentage, estimated by mapping 150,000 unique merged reads > 30bp to the polar bear reference genome assembly. Estimates obtained using the Dabney DNA isolation and single-stranded library protocols are shown.

A schematic of the experimental design is shown in Figure 1. All laboratory work preceding PCR amplification was carried out in dedicated ancient DNA facilities at the University of Potsdam, following established procedures to prevent contamination with modern or synthetic DNA (Fulton 2012). Negative controls were included in all experiments. 350 mg of bone powder from each petrous bone was digested overnight in 7ml extraction buffer (0.45M EDTA, 0.25mg/ml Proteinase K). The resulting supernatant was then divided into three equal 2ml aliquots and each subjected to a different DNA isolation protocol, producing 50μl final DNA extract each. These DNA extracts were then each divided into two equal 20μl aliquots for conversion into Illumina sequencing libraries using either a double-or single-stranded method.

**Figure 1.**
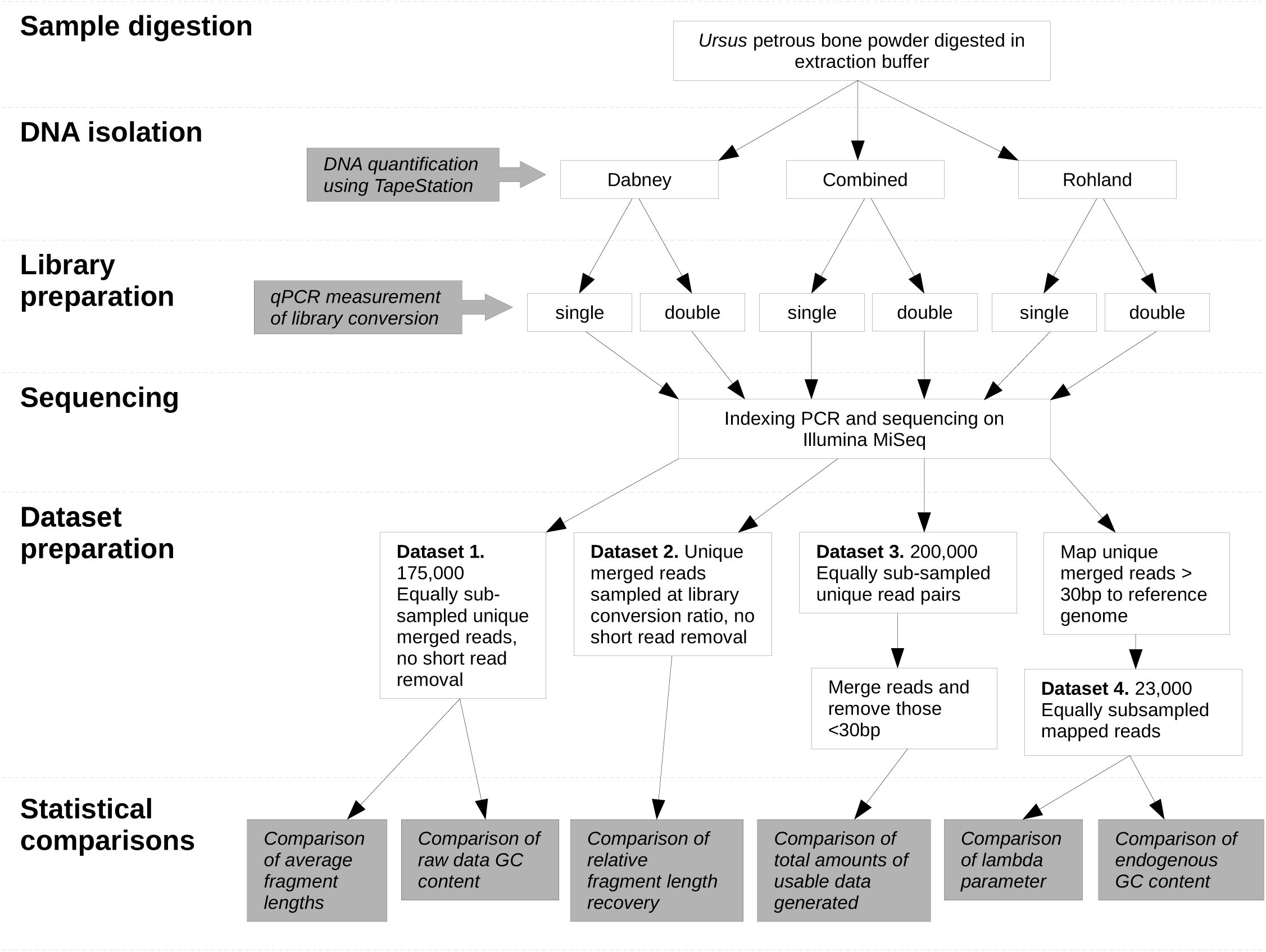
Hierarchical experimental design for testing of three alternative DNA isolation protocols and two alternative library protocols, and their interaction. Stages of investigation are indicated to the left of the figure. Grey boxes indicate specific comparisons presented in the results.

The DNA isolation protocols tested were that of Dabney *et al*. (2013) (“Dabney protocol”) and that of Rohland *et al.* (2010) (“Rohland protocol”), in addition to a third novel protocol (“combined protocol”). The Dabney protocol is a modified version of a standard commercial spin-column (Qiagen) DNA isolation method, involving the binding DNA to a silica membrane in the presence of a binding buffer containing guanidine hydrochloride, sodium acetate and isopropanol. The Dabney protocol uses a large ratio of binding buffer to sample (13:1) to maximise DNA recovery. Our procedure followed exactly that described in Dabney *et al.* (2013), except that 2ml supernatant from the bone digest was mixed with 26ml binding buffer and the resulting mixture passed through a single spin-column in two equal aliquots. The Rohland protocol is similar in principle except that DNA is bound to silica particles in suspension, a binding buffer containing only guanidinium thiocyanate and sodium acetate is utilised, and that the ratio of sample to binding buffer is lower than that of the Dabney protocol. Our procedure followed exactly that described in Rohland *et al.* (2010), except that 2ml supernatant from the bone digest was mixed with 1ml binding buffer and 40μl silica suspension. The combined protocol was designed to capture the key components of both of these protocols. Using the approach of the Rohland protocol, DNA molecules were bound to silica particles. The ratio of sample to binding buffer was then increased to match that of the Dabney protocol, and the resulting mixture passed through a silica spin column to capture any remaining DNA not bound to the silica particles. A full account of the combined protocol is provided in the supporting information (Appendix S1, Supporting Tables S1 and S2). DNA extracts were quantified using an Agilent 2200 TapeStation instrument with D1000 High Sensitivity ScreenTape and reagents.

DNA extracts resulting from each of the three purification protocols were then converted into sequencing libraries using either the double-stranded approach described in Meyer & Kircher (2010), with the modifications described in Fortes & Paijmans (2015), or the single-stranded approach described in Gansauge & Meyer (2013). The double-stranded library protocol involves the ligation of double-stranded Illumina P5 and P7 adapters to each end of blunt-ended DNA molecules, respectively. As the ligation is random, about half of the template molecules will receive incorrect combinations of adapters and will be lost from the final library after amplification. The double-stranded library protocol also requires a number of purification steps each of which is associated with the loss of molecules from the remaining library. The single-stranded library protocol, in contrast, denatures double-stranded DNA into its component single strands, and then ligates a biotinylated adapter oligo to the 3′ end of each molecule. After immobilisation on Streptavidin-coated beads, a complementary oligo is annealed to the adapter and the complementary template DNA strand filled in, after which the P5 adapter can be ligated to the free end of the template molecule. Prior to adapter ligation, DNA extracts processed using the single-stranded library protocol were treated with the enzymes uracil-DNA glycosylase (UDG) and endonuclease VIII to remove deoxyuracils resulting from postmortem DNA damage.

The efficiency of library conversion was investigated using qPCR assay of unamplified libraries with primers complementary to the P5 and P7 adapter sequences flanking the DNA insert. Four replicate 10μl reactions were carried out with amounts of template corresponding to 0.2% of the total library. All other experimental details followed those described in Gansauge & Meyer (2013). Mean Ct values across all replicates were calculated, except where single replicates differed from this mean by ±0.5 Ct, which were discarded and a new mean calculated.

A unique index sequence was incorporated into the P7 adapter of each library during library amplification. Indexing of single-stranded libraries utilised AccuPrime Pfx DNA polymerase (ThermoFisher) and was carried out in 4 parallel 20μl reactions. Indexing of double-stranded libraries utilised the Accuprime SuperMix 1 PCR mastermix, for its ability to read over uracils, and was carried out in four parallel 25μl reactions. The number of amplification cycles was varied according to the results of qPCR experiments (Gansauge & Meyer 2013). Indexed libraries were pooled in an equimolar ratio and sequenced on a single flowcell of the Illumina MiSeq sequencing platform, producing 2x70bp paired-end (PE) reads.

### Sequence data processing

Processing of sequence reads involved removal of duplicate sequences using FastUniq (Xu *et al.* 2012) and trimming of adapter sequences using CutAdapt (Martin 2011). Overlapping paired-end reads were merged using the program Flash (Magoč & Salzberg 2011).

Sequencing of single-stranded libraries requires a custom R1 sequencing primer, which, when applied to double-stranded libraries, results in 5bp of the P5 adapter sequence at the 5′ end of read 1. These 5bp were removed, also using CutAdapt, reducing the maximum length of merged reads from double-stranded libraries from 130bp to 125bp. Merged reads from single-stranded libraries >125bp in length were also discarded to allow unbiased comparisons of read lengths. To avoid biases resulting from the unequal sampling of reads, the trimmed and merged reads were randomly subsampled using the program SeqTK (Li 2012). To separate endogenous reads from those of exogenous (contaminant) sources, we mapped reads to the genome assembly of the closely related polar bear (Li *et al.* 2011) using the program bwa (Li & Durbin 2009) with the default mismatch parameter and processed the resulting alignment using SAMtools (Li *et al.* 2009). Only merged reads were used so that the length of the original template molecules could be determined. Merged reads <30bp were not used for mapping to reduce the probability of spurious alignment.

### Statistical comparisons of DNA isolation and library protocols

The effect of different protocols, and their interaction, was assessed by investigating five different data properties: 1. total DNA recovery, 2. fragment length recovery, 3. lambda, 4. GC content, and 5. total amount of usable data generated. Sequence-based comparisons were made using four sets of data (Datasets 1-4), which are described in Figure 1.

#### 1. Total DNA recovery

Rates of DNA recovery of different DNA isolation protocols were compared using the results of TapeStation analysis of DNA extracts.

The rate of conversion of ancient DNA molecules into sequencing libraries by the single-and double-stranded protocols was compared using their mean CT values obtained in the qPCR experiment. The relative difference in the numbers of molecules converted into library (conversion ratio) was then calculated by the formula:

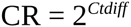

Where CR is the conversion ratio and *Ctdiff* is the difference in the number of cycles required to reach a particular amplification level in the double-stranded library minus that required for the single-stranded library. CR=1 when the number of molecules converted into each library is equal. CR>1 when single-stranded conversion exceeds double-stranded conversion, while CR<1 when double-stranded conversion exceeds single-stranded conversion. This model assumes perfect PCR amplification efficiency.

#### 2. Fragment length recovery

The effect of different DNA isolation and library protocols, and their interaction, on average fragment length recovery was investigated using Dataset 1 (Fig. 1). We additionally investigated the mechanism by which alternative library protocols produced different average fragment lengths by re-sampling reads in proportions equal to their ratios of conversion (Dataset 2, see Fig. 1). This was accomplished through keeping the number of reads from single-stranded libraries at 175,000 and adjusting the number of reads sampled from double-stranded libraries accordingly.

#### 3. Lambda

The effect of different DNA isolation and library protocols, and their interaction, on estimates of the lambda DNA degradation parameter was investigated using Dataset 4 (Fig. 1). Lambda was calculated by linear regression of log-transformed fragment frequencies between 70– 100bp. Lambda values were not considered if log-transformed fragment length distributions showed a poor fit to the linear model (adjusted *R*2 < 0.8). This approach limited statistical comparisons to only those values obtained from libraries generated using the single stranded protocol.

#### 4. GC content

The effect of different DNA isolation and library protocols, and their interaction, on GC content of the raw sequencing data (unmapped reads, i.e. prior to any mapping being carried out), was investigated using Dataset 1 (Fig. 1). We also investigated effects on the GC content of endogenous sequence data (mapped reads) using Dataset 4 (Fig. 1).

#### 5. Total amount of usable data generated

The effect of different DNA isolation and library protocols, and their interaction, on the total amount of usable data generated was deduced from Dataset 3 (Fig. 1). Comparisons were based on the proportion of read pairs within each sample treatment that were 1) < 30bp after adapter trimming and merging, and thus deemed too short for reliable mapping; 2) Merged > 30bp; and 3) Unmerged, having insufficient overlap. We additionally compared the total amounts of sequence data (in number of bases) provided by the merged reads and by the merged reads plus unmerged reads. The latter comparisons were log-transformed prior to significance testing.

### Significance testing

For sequence-based dependent variables, significant effects of different protocols were assessed using linear mixed-effects model analysis, following the procedures described by Winter (2013). Linear mixed-effects models are a highly flexible class of statistical models that allow the effects of a number of experimental variables (fixed effects) to be assessed simultaneously, while accounting for variation resulting from additional factors (random effects) that may also influence the results. These methods are appropriate in our case because they accommodate the hierarchical nature of the experimental design (Fig. 1) and provide a means of accounting for inter-sample variability, which may be substantial among subfossil bone samples due to variable ages and deposition microenvironments. For analysis of each dependent variable, DNA isolation and library protocol, and their interaction, were assigned as fixed effects. Sample was assigned as a random effect with different random intercepts for each of the four bone samples and random slopes for DNA isolation and library protocol. The significance of each fixed effect was then assessed by comparing the likelihood of this full model against a null model with the fixed effect in question removed, using likelihood ratio test. Analyses were performed in R (R Core Team 2014) using the lme4 (Bates *et al.*2014) and Effects (Fox 2009) packages, with appropriate checks for deviations from assumptions of homoscedasticity and normality.

## RESULTS

All data used for used for statistical analysis are provided in Supporting Table S3.

### Total DNA recovery

The amount of DNA recovered by different isolation protocols varied considerably (Fig. 2). The Dabney protocol consistently recovered more DNA than both the Rohland and the combined protocol. On average, the Dabney protocol provided a 14.29-fold (minimum 1.01-fold, maximum 42.95-fold) increase in DNA recovery over the combined protocol, and a 12.60-fold increase over the Rohland protocol (minimum 4.3-fold, maximum 35.89-fold).

**Figure 2.**
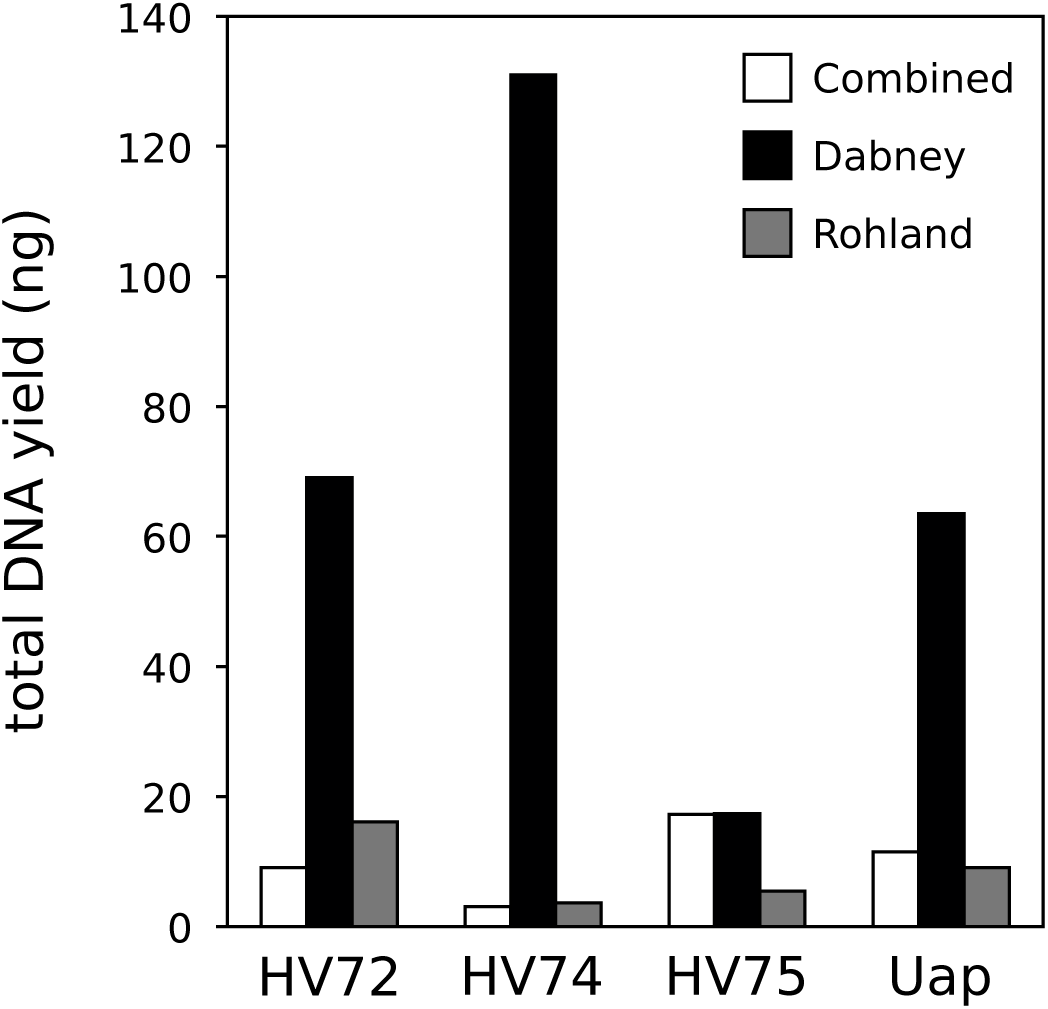
Total DNA obtained from four ancient bone samples using three DNA isolation protocols: combined (open bars), Dabney (black bars) and Rohland (grey bars). Each DNA extract derived from the equivalent of 100mg bone powder and was eluted in 50μl volume. DNA was quantified using an Agilent TapeStation instrument within a 30–1000 bp range.

Substantial differences in DNA recovery were also found between the single-and double-stranded library protocols (Fig. 3). The single-stranded library protocol converted more molecules than the double-stranded protocol in seven out of nine library comparisons. In the two cases where conversion was lower using the single-stranded method, the difference was minor, with a maximum 1.3-fold increase in conversion for the double-stranded protocol. In contrast, in comparisons where the single-stranded method performed best, the difference was as much as 122-fold, with a mean increase of 25.6-fold. This value is greatly influenced by two comparisons with very high conversion rates (122-and 32-fold), but even when these are excluded the mean increase in conversion remains substantial, at 4.6-fold.

**Figure 3.**
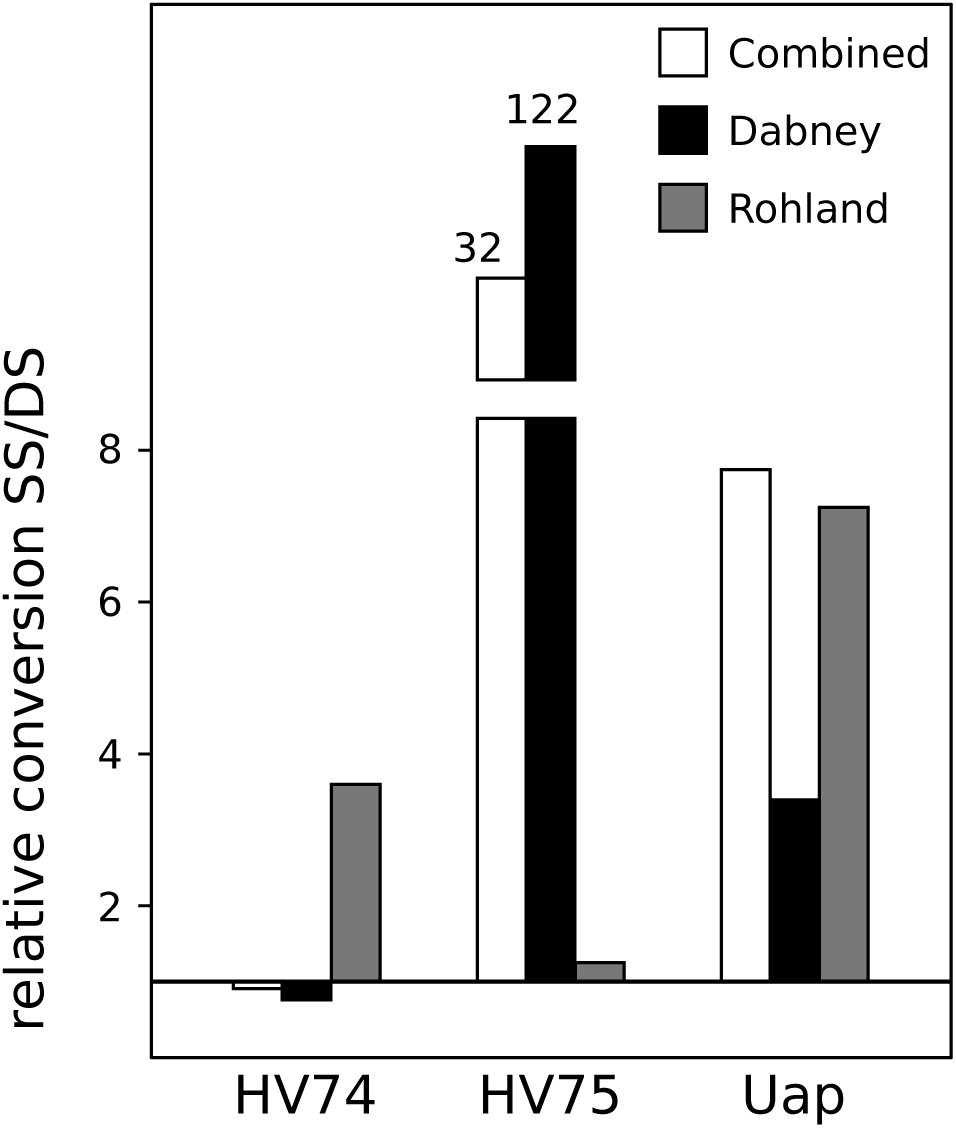
Relative conversion rates of single-and double-stranded library protocols applied to nine DNA extracts. DNA isolation protocol used to obtain these extracts is indicated following the colour coding shown in Figure 1. The horizontal line intersecting the y-axis at 1 indicates equal conversion. Note that for two extracts obtained from sample HV74, single-stranded conversion was lower than double-stranded. Also, conversion was dramatically higher using the single-stranded protocol for two extracts obtained from sample HV75. These are shown as broken bars, with the actual conversion ratio indicated above.

### Fragment length recovery

We found significant effects of both DNA isolation and library protocol on the length of recovered ancient DNA fragments, in addition to a significant interaction effect of these factors (Table 2). The single-stranded library protocol consistently produced shorter mean fragment lengths than the double-stranded protocol, irrespective of DNA isolation method (Fig. 4). Among DNA isolation protocols, the Rohland protocol consistently recovered longer fragment lengths than either the combined or the Dabney protocol, with this difference been considerably larger when the double-stranded library method is utilised. Differences between the combined and Dabney protocols were less pronounced and inconsistent between library protocols (Fig. 4). Analysis of median fragment lengths also showed significant effects of both factors with a significant interaction (Table 2), with comparable changes as found for mean lengths (Supporting Fig. S1).

**Figure 4.**
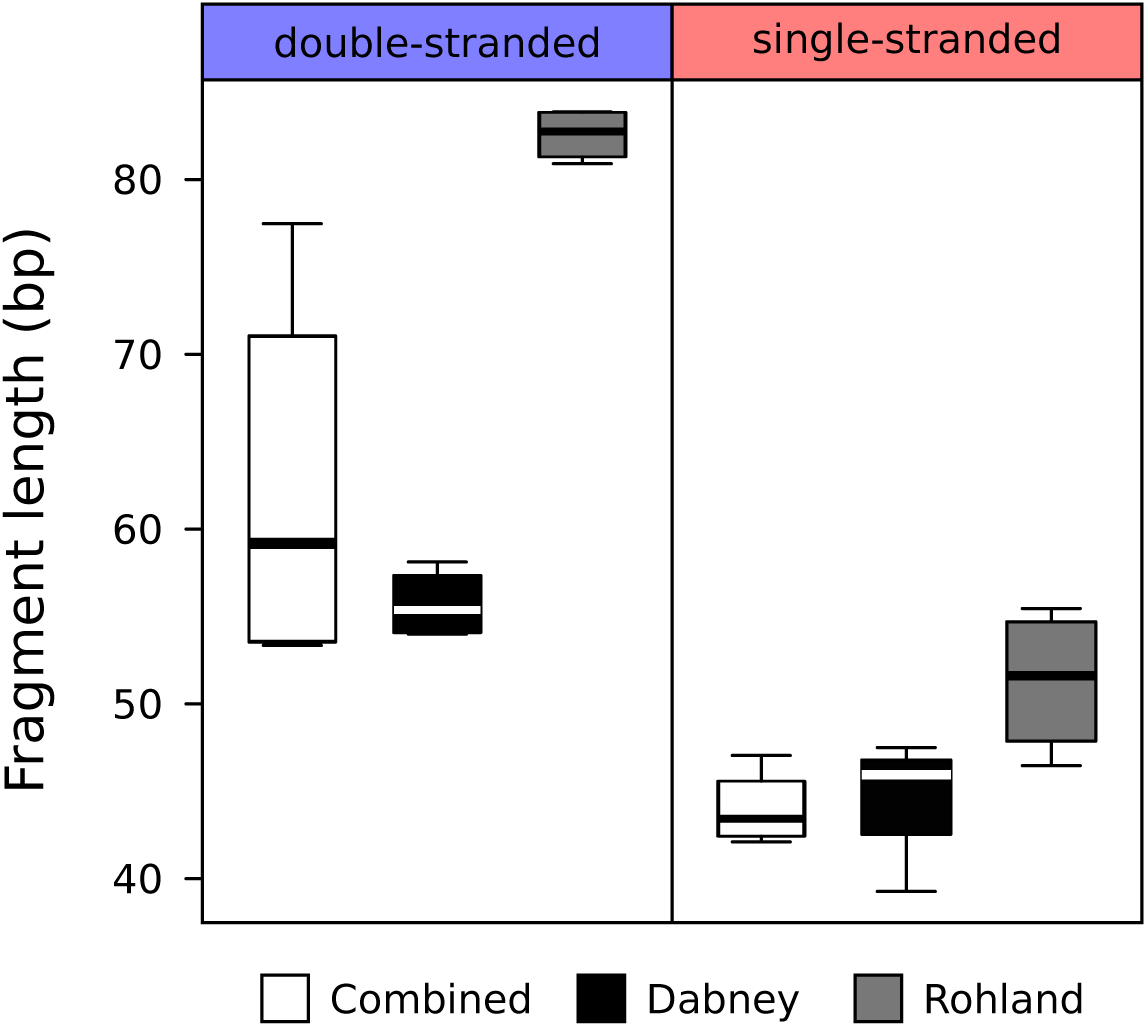
Boxplots showing the effect of different DNA isolation and library protocols on total mean fragment length recovery from four ancient bone samples. Library protocol is indicated above boxplots, and DNA isolation protocol below.

**Table 2.** Significance of effect of DNA extraction and library protocol on palaeogenomic data.

| **Data Property** | **Factor** | **Chi2** | **df** | **P-value\*** |
| --- | --- | --- | --- | --- |
| mean fragment length | DNA extraction | 37.362 | 4 | < **0.01** |
|  | Library | 37.505 | 3 | < **0.01** |
|  | DNA extraction * Library | 26.218 | 2 | < **0.01** |
| median fragment length | DNA extraction | 42.070 | 4 | < **0.01** |
|  | Library | 43.490 | 3 | < **0.01** |
|  | DNA extraction * Library | 32.368 | 2 | < **0.01** |
| Lambda | DNA extraction | 15.783 | 2 | < **0.01** |
| GC content unmapped reads | DNA extraction | 7.808 | 4 | 0.099 |
|  | Library | 15.350 | 3 | < **0.01** |
|  | DNA extraction * Library | 5.970 | 2 | 0.051 |
| GC content mapped reads | DNA extraction | 26.252 | 4 | < **0.01** |
|  | Library | 31.253 | 3 | < **0.01** |
|  | DNA extraction * Library | 10.385 | 2 | < **0.01** |
| short reads <30bp | DNA extraction | 8.451 | 4 | 0.076 |
|  | Library | 22.669 | 3 | < **0.01** |
|  | DNA extraction * Library | 3.739 | 2 | 0.154 |
| merged reads | DNA extraction | 34.790 | 4 | < **0.01** |
|  | Library | 29.785 | 3 | < **0.01** |
|  | DNA extraction * Library | 26.312 | 2 | < **0.01** |
| unmerged PE reads | DNA extraction | 34.030 | 4 | < **0.01** |
|  | Library | 30.388 | 3 | < **0.01** |
|  | DNA extraction * Library | 22.857 | 2 | < **0.01** |
| total merged bp | DNA extraction | 8.467 | 4 | 0.076 |
|  | Library | 17.336 | 3 | < **0.01** |
|  | DNA extraction * Library | 4.699 | 2 | 0.095 |
| total bp (merged + unmerged PE) | DNA extraction | 23.962 | 4 | < **0.01** |
|  | Library | 21.443 | 3 | < **0.01** |
|  | DNA extraction * Library | 10.085 | 2 | < **0.01** |

\*P-values below the 0.05 threshold of statistical significance are indicated in bold.

The reduction in average length of recovered fragments in the single-stranded library protocol may potentially result from either a relative increase in short fragment recovery or, conversely, a relative decrease in long fragment recovery. These alternative mechanisms were investigated using fragment length distributions of single-and double-stranded libraries sampled at their conversion ratios (Fig. 5). Five out of nine comparisons indicate either no reduction or an increase in long fragment recovery for the single-stranded protocol (Figs. 5a, 5b, 5c, 5e, 5f). This suggests that the reduction in average fragment length observed for the single-stranded protocol results from an increase in short fragment recovery rather than a loss of long fragments.

**Figure 5.**
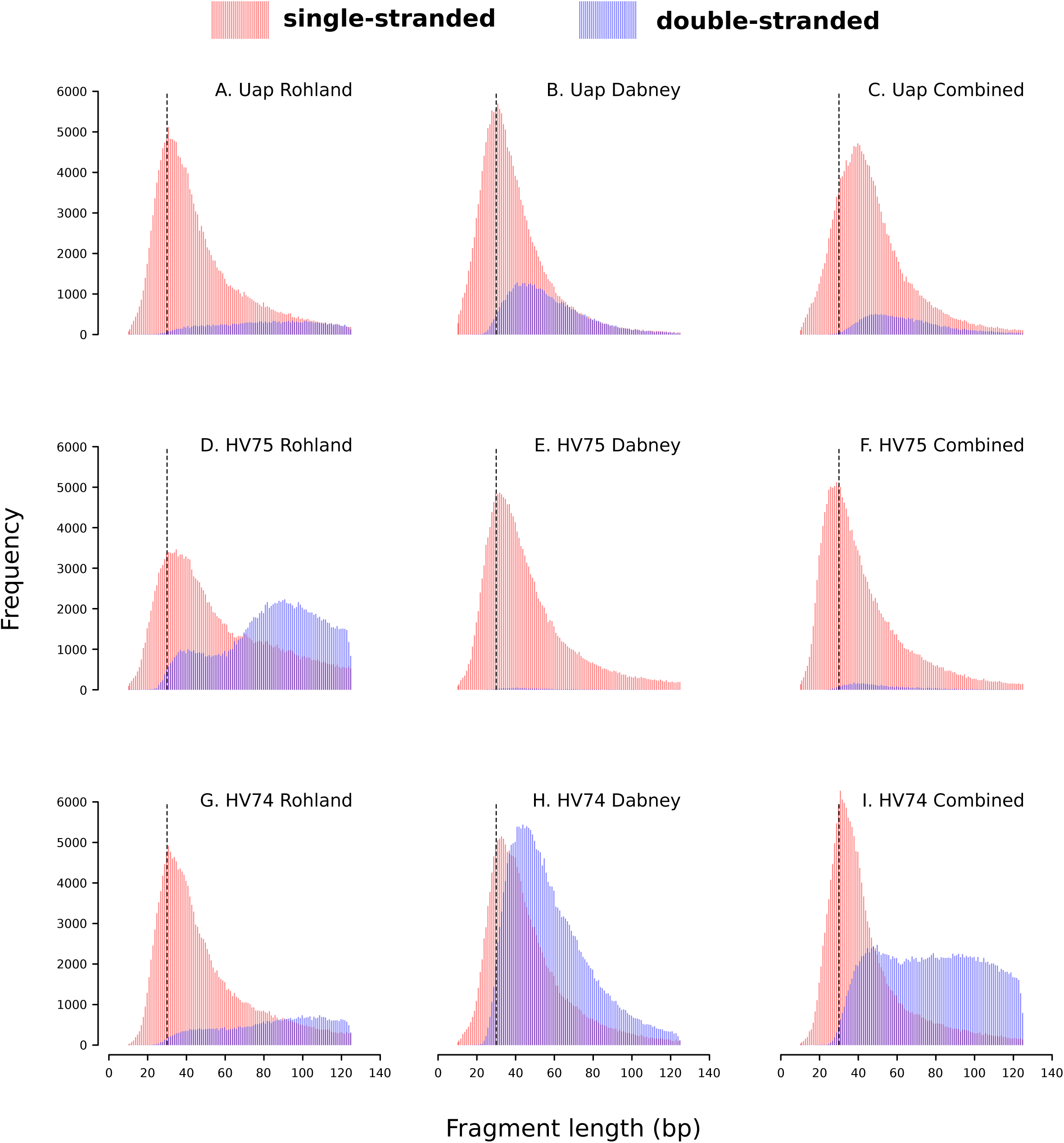
Length distributions of total fragments obtained using the single-stranded (red) and the double-stranded (blue) library protocol applied to the same DNA extract. Sample and DNA extraction protocol are indicated above each comparison. Sampling of reads for each library pair is scaled according to their conversion ratio estimated using qPCR (see Fig. 2). Vertical dashed lines indicates a 30bp minimum read length threshold frequently employed by ancient DNA, where reads shorter than this threshold are discarded prior to mapping.

### Lambda

We were unable to generate meaningful lambda values for any libraries generated using the Rohland DNA isolation protocol in conjunction with the double-stranded library protocol (adjusted *R*2 of log-transformed read lengths < 0.8), as well as for one library generated using the combined DNA isolation protocol and double-stranded library protocol (sample HV74).

Examination of these read length distributions (Fig. 5) show they do not conform to the expected pattern for ancient DNA of exponential reduction in frequency of fragments longer than the mode. This observation alone strongly suggests a significant effect of both DNA isolation and library protocol, in addition to a significant interaction effect, on lambda estimates. This hypothesis is further supported by statistical analysis of libraries generated using the single-stranded library protocol, which confirm a significant effect of DNA isolation protocol on lambda (Table 2). The ranked order of decreasing lambda is Dabney – combined – Rohland, with the greatest shift in lambda observed between the combined and Rohland protocols (Fig 6).

**Figure 6.**
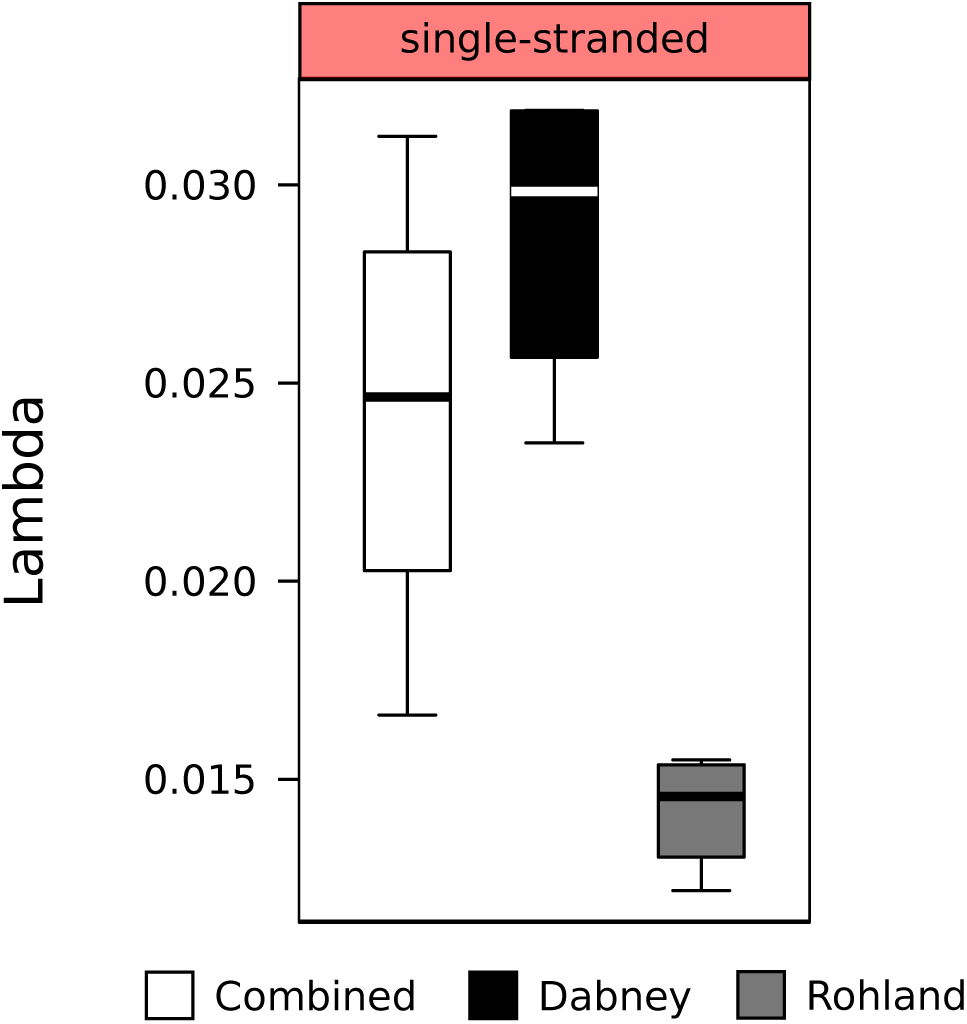
Boxplots showing the effect of different DNA isolation and library protocols on lambda DNA degradation parameter estimates for endogenous DNA molecules from four ancient bone samples. Results are based only on libraries generated using the single-stranded library protocol. DNA isolation protocol is indicated below boxplots.

### GC content

A significant effect of library protocol, but not of DNA isolation protocol nor any interaction effect, were found on the GC content of unmapped reads (Table 2): double-stranded libraries resulted in higher GC contents than single-stranded libraries, irrespective of the DNA isolation protocol applied (Fig. 7). We did observe substantial inter-sample variation in GC content, however, and so with increased sampling it may be that a significant effect of DNA isolation protocol is detectable.

**Figure 7.**
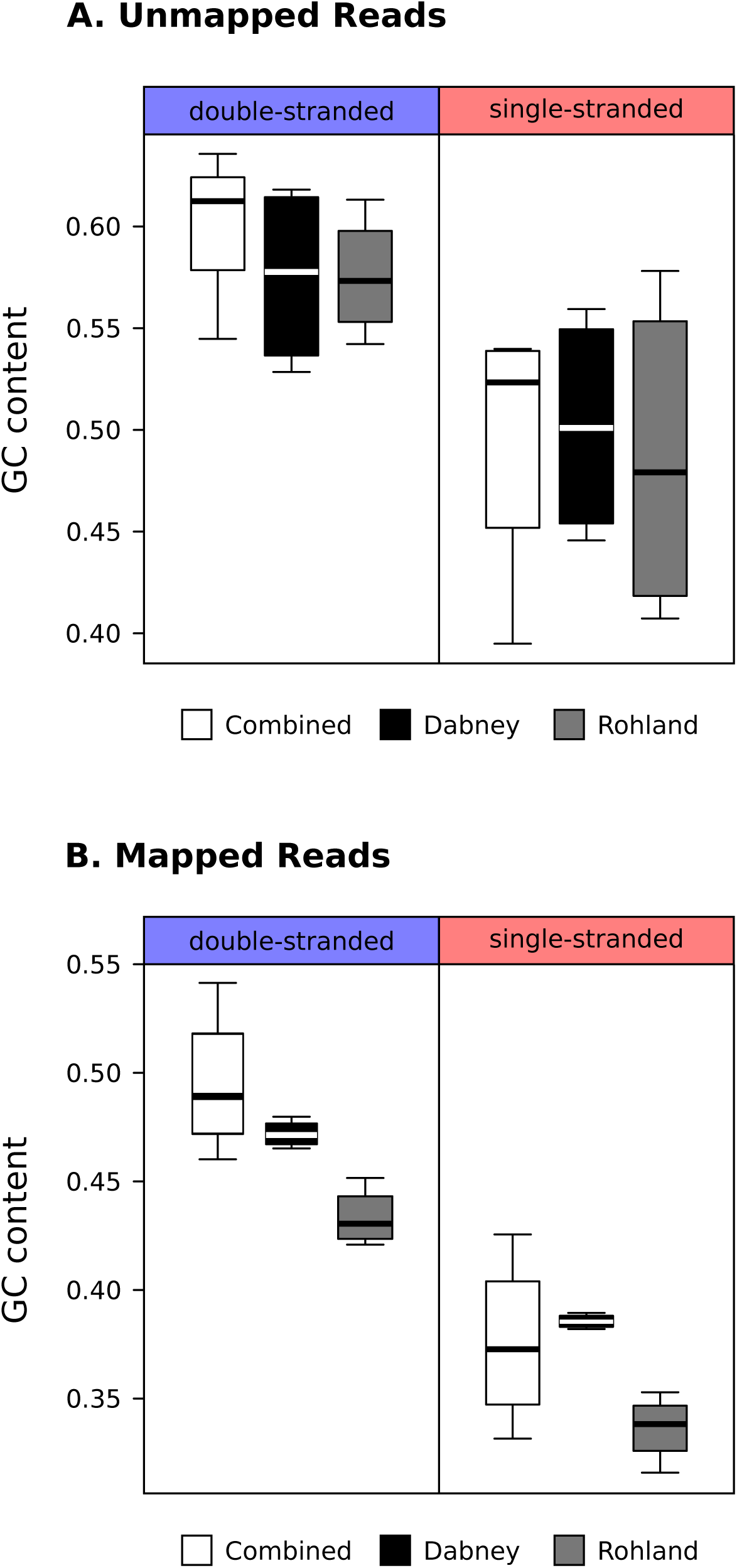
Boxplots showing the effect of different DNA isolation and library protocols on data GC content for four ancient bone samples, for A) total (unmapped) reads and B) endogenous (mapped) reads. Library protocol is indicated above boxplots, and DNA isolation protocol below.

In contrast to the unmapped reads, analysis of mapped reads did support significant effects of both DNA isolation and library protocol, in addition to a significant interaction effect, on GC content (Table 2). Again, the highest GC contents were observed using the double-stranded library protocol (Fig. 7). Within each library type, the Rohland DNA isolation protocol produced the lowest GC contents. As for the Dabney and combined DNA isolation protocols, the ranked order of mean GC content was variable among library types. It is also notable that the inter-sample variability in GC content was dramatically reduced when the Dabney DNA isolation protocol was employed both for the single-and double-stranded library protocols, although with higher overall GC content in the latter case.

### Total amount of usable data generated

Analysis of read proportions revealed that DNA isolation protocol, library protocol, and their interaction, have significant effects on the proportions of merged and of unmerged reads generated (Table 1). Library protocol also had a significant effect on the proportion of reads < 30bp, but no significant effect of DNA isolation, nor any significant interaction, was found (Table 1). In general, the single-stranded library protocol produced a substantially greater proportion of reads < 30bp than the double-stranded protocol. The Rohland DNA isolation protocol produced a greater proportion of unmerged read-pairs than either the combined or the Dabney protocols, with the greatest increase being associated with the double-stranded library protocol (Fig. 8).

**Figure 8.**
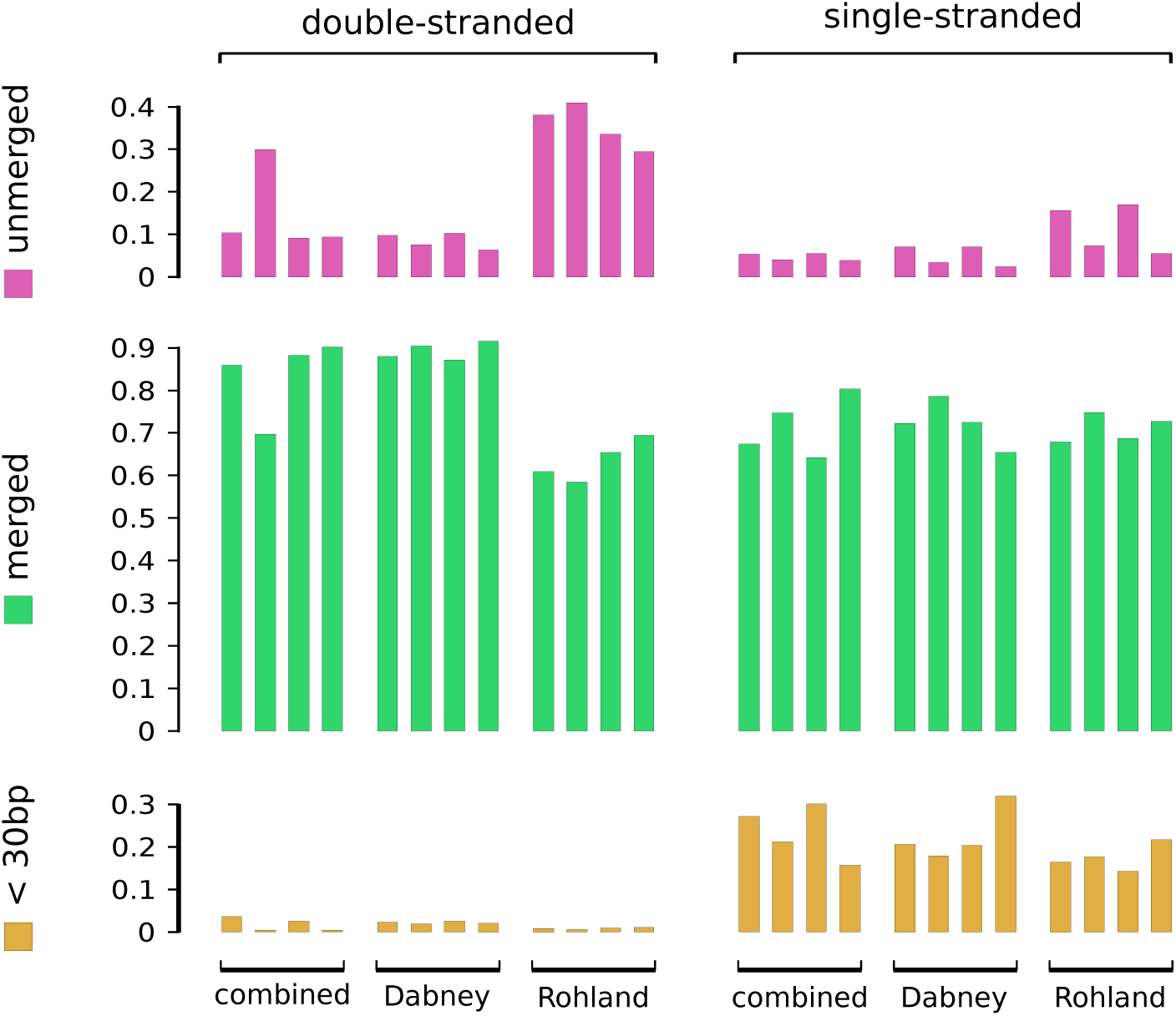
Effect of different DNA isolation and library protocols on the total proportion of short (< 30bp), merged, and unmerged, sequence reads. Y axes indicate the proportion of reads in each category. Vertically aligned sets of bars correspond to a particular sample, with DNA isolated using one of three alternative protocols (indicated below bars), and converted into sequencing libraries using one of two alternative protocols (indicated above bars). Within each treatment group, vertical sets are ordered from left to right corresponding to samples HV72, HV74, HV75, and Uap.

Similarly, DNA isolation protocol, library protocol, and their interaction, were found to have significant effects on the total amount of usable sequence data (merged and unmerged bases of reads >30bp) generated (Table 1). The double-stranded library protocol consistently produced more usable data than the single-stranded protocol, when applied to the same DNA extract (Fig. 9). Among DNA isolation methods, the Rohland protocol produced substantially more total usable data than either the Dabney or the combined protocol, again with the greatest increase being associated with the double stranded library protocol (Fig. 9). Only library protocol, and not DNA extraction or their interaction, was found to have a significant effect of the total amount of merged data, with larger amounts of data generated by the double-stranded protocol (Fig. 9; Table 1).

**Figure 9.**
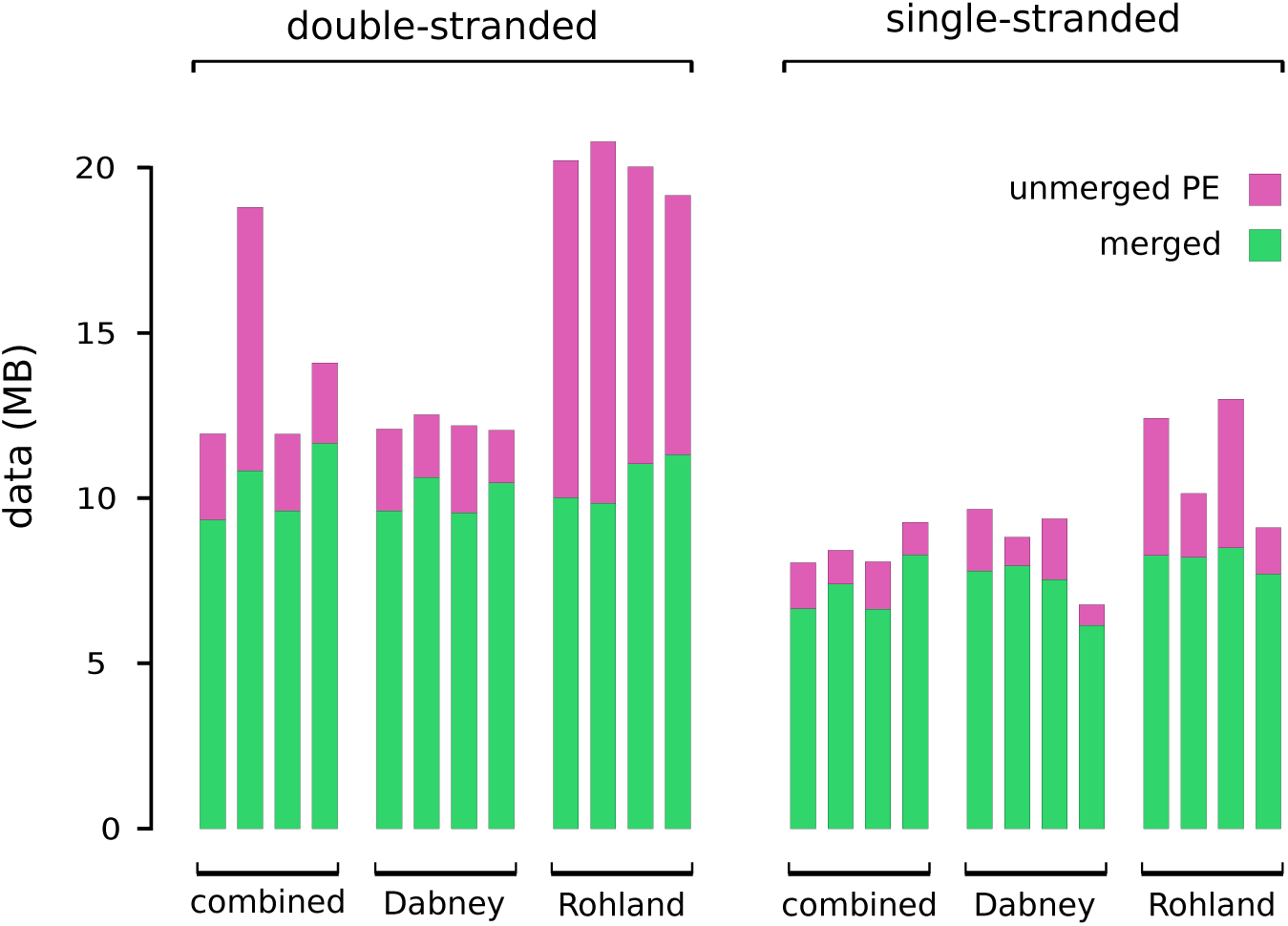
Effect of different DNA isolation and library protocols on the total yield of usable sequence data, in MB (Y axis), from a total of 28MB of sequence data. Each bar corresponds to a particular sample, with DNA isolated using one of three alternative protocols (indicated below bars), and converted into sequencing libraries using one of two alternative protocols (indicated above bars). Within each treatment group, vertical sets are ordered from left to right corresponding to samples HV72, HV74, HV75, and Uap. Bars have been divided according to the proportion of sequence data derived from merged (green) and unmerged (pink) read pairs.

## DISCUSSION

Our results show that palaeogenomic data obtained from ancient subfossil bone samples is greatly influenced by the methods of DNA isolation and library preparation that are utilised. These effects include changes in the lengths of recovered fragments, nucleotide composition, and total data yields. Our results have widespread implications for palaeogenomic research. Laboratory methods not only directly influence the quantity of data that can be recovered from ancient samples, but may also be sufficient to confound analysis and interpretation of the data.

### Effects on fragment length recovery

Ancient DNA is short. This fact has motivated the development of protocols that enhance short fragment recovery. It is therefore unsurprising that we found significant changes in the average length of fragments recovered using different DNA isolation and library preparation protocols. In agreement with previous studies (Meyer *et al.* 2012; Gamba *et al.* 2016), we found the average fragment length recovery of the Dabney DNA isolation protocol to be considerably shorter than that of the Rohland protocol, but our novel combined protocol also proved to be capable of recovering similarly short fragments (Fig. 4). Comparison of library protocols showed, in agreement with previous studies (Meyer *et al.* 2012; Bennett *et al.* 2014; Wales *et al.* 2015), mean fragment length recovery of the single-stranded library protocol to be shorter that of the double-stranded protocol (Fig. 4). We additionally found a significant interaction effect between DNA isolation and library preparation protocol. For instance, the increase in average fragment length associated with the Rohland DNA isolation protocol is greatly enhanced when used in conjunction with the double-stranded, relative to single-stranded library protocol. This result demonstrates the importance of considering the entire suite of laboratory methods applied to a sample in predicting effects on the resulting data.

The observed size distribution of ancient DNA can be used as a method of verifying its authenticity (Noonan *et al.* 2006; Ginolhac *et al.* 2011). However, employing laboratory methods that modify the fragment length distribution could lead to a degree of circularity in this argument. Since all methods likely provide some level of fragment length bias, the true distribution of fragment lengths in a bone sample is difficult to ascertain. Our data do provide information on relative shifts, however, which proved dramatic in some cases. In particular, use of the Rohland, and in one instance the combined, DNA isolation protocol in conjunction with the double-stranded library protocol produced read length distributions that deviated substantially from that expected for ancient DNA. Overall, we found the Dabney DNA isolation protocol used in conjunction with the single-stranded library protocol consistently provided fragment length distributions that conformed to expectations, and may therefore represent an optimal set of methods for assessments of data authenticity.

An extension of the qualitative comparison of fragment length distributions is the calculation of lambda: the rate of decay of fragment length frequencies. Lambda estimates have been used to compare DNA degradation among ancient samples as a product of sample age and the thermal deposition environment (Allentoft *et al.* 2012). Our results clearly show that in addition to these environmental factors, the method of DNA isolation and library preparation can also significantly and substantially influence lambda estimates, even to the extent that no accurate estimate of lambda can be made. We therefore conclude that any experimental investigation of ancient DNA fragmentation rates will need to carefully consider the laboratory methods used to generate empirical measures, which may need statistical correction to account for any biases that are introduced.

### Effects on nucleotide composition

Raw, unmapped data obtained from ancient bone samples represents a metagenomic dataset of the target species and associated contaminating organisms. It is unsurprising that we observed large variation in the GC content of unmapped reads among samples, since the biota in their respective deposition microenvironments is likely to vary substantially (Der Sarkissian *et al.* 2014). However, we additionally detect a significant effect of library preparation on GC content of the unmapped reads, with the single-stranded method showing an overall reduction in comparison to the double-stranded method, which has also been observed previously (Wales *et al.* 2015). It is noteworthy that the single-stranded libraries were treated with UDG/EndoVIII to remove uracils prior to library preparation, whereas double-stranded libraries were not. However, omission of UDG/EndoVIII treatment would lead to a reduction in overall data GC content of double-stranded libraries relative to single-stranded, which is the opposite of what we find. Thus, our results likely underestimate the magnitude of the difference in GC content associated with these library protocols. The biases in GC content introduced at the stage of library preparation is of particular relevance for the field of metagenomic and environmental ancient DNA analysis (e.g. Willerslev et al. 2007; Der Sarkissian et al. 2014), and future studies applying these methodologies may be advised to consider whether such biases may affect estimates of organismal diversity and abundance.

For studies of targeted species, the nucleotide composition of reads mapped to a reference genome is likely to be of greater relevance. Here, we find significant effects of both DNA isolation and library protocol, as well as a significant interaction effect, on GC content. In line with our results, relative increases in GC content have been reported previously for double-in comparison to single-stranded library preparations (Wales *et al.* 2015). Changes in GC content have similarly been reported for the Rohland DNA isolation protocol in comparison the Dabney protocol (Gamba *et al.* 2014), but in the opposite direction to that observed for our samples. This is likely explained by sample-dependent effects, which reinforces the difficulty of making generalised predictions based on data from and small number of samples. Shifts in library GC content have also been demonstrated for some DNA polymerases during successive cycles of library amplification (Dabney & Meyer 2012). However, it is highly unlikely that this is the primary factor explaining our results as comparisons of GC content and amplification cycle showed no obvious correlation (Supporting Figure S2). Rather, substantial GC content bias appears to be an inherent property of some of the DNA isolation and library preparation protocols investigated here.

It is reasonable to assume that all cave bears had similar average genomic GC contents, and that cave bear GC content is broadly similar to those reported from genome assemblies of other representatives of the Carnivora (e.g. *Ursus maritimus* 43.2%, *Felis catus* 40.6%, *Canis familiaris* 40.8%). However, by applying different laboratory methods, we were able to obtain completely unrealistic GC contents ranging from 31.6% to 54.1% for mapped endogenous data from a single sample (Uap, values for Rohland + single-stranded, and combined + double-stranded, respectively), and mean values across all samples that range from 33.6% to 49.5% (Fig. 7b: Rohland + single-stranded, combined + double-stranded, respectively). Moreover, most combinations of DNA isolation and library protocol produced GC contents that were highly variable among samples (see large interquartile and maximum/minimum ranges in Fig. 7b). Thus, empirical GC contents of the mapped reads cannot be explained in terms of a simple, fixed upward or downward bias in nucleotide composition. The exceptions to this pattern were datasets generated using the Dabney DNA isolation protocol, which produced highly consistent GC contents among samples, with the most realistic values obtained when used in conjunction with the single-stranded library protocol.

In addition to distorting overall measures of genomic GC content, it is feasible that biases in nucleotide composition of the magnitude detected here could also confound identification of DNA polymorphisms and measures of genetic diversity, as well as estimates of mutation rates. These effects would likely be exacerbated at lower levels of genomic coverage, which have been frequently utilised by palaeogenomic studies (Leonardi *et al.* 2016), and in mixed datasets where different methods are applied across samples. The latter situation is likely to become increasingly important as the number of published paleogenomics datasets continues to grow, leading to combined analyses of multiple datasets from multiple modern and ancient samples, generated using a variety of alternative laboratory methods.

### Efficiency of data production

Previous comparisons of laboratory methods for palaeogenomic studies have generally focussed on identifying methods that improve the cost-or time-efficiency of data production (Bennett et al. 2014; Wales *et al.* 2015; Gamba et al. 2016). Although other factors, such as obtaining unbiased and representative sampling of the genome, may ultimately be of greater importance, the efficiency of data production can impose practical limits on palaeogenomic studies and is therefore worthy of discussion. Palaeogenomic data production efficiency is principally determined by: 1) sample complexity, which is simply the total number of unique molecules contained within a DNA extract or library; and 2) the total amount of usable sequence generated per unit of sequencing effort, which is determined by the proportion of endogenous DNA in the sample as well as the length of recovered endogenous fragments relative to sequencing read length. Our results show that the methods of DNA isolation and library preparation can have substantial effects on data production efficiency by influencing both these factors.

The effect of reduced sample complexity on sequencing data is increased read duplication, which scales exponentially relative to sequencing effort until all molecules in the library have been sequenced. Thus, any improvement in DNA recovery at the stage of either DNA isolation or library preparation will reduce duplication rates and improve data production efficiency, particularly when very low quantities of DNA are present, which can be considered typical for ancient samples (Hofreiter *et al.* 2014). Quantification of our cave bear DNA extracts showed the Dabney protocol to recover, on average, an order of magnitude more total DNA than either the Rohland or combined DNA isolation protocols. A previous study similarly reported increased yields using the Dabney protocol versus the Rohland protocol from indirect measures of data duplication rates (Gamba *et al.* 2016). Among library preparation protocols, qPCR measurement of conversion rates showed that the single-stranded protocol typically led to much higher levels of conversion than the double-stranded method, which has also been reported previously (Gansauge & Meyer 2013; Wales *et al.* 2015). Examination of read length distributions scaled to conversion rates indicates that the enhanced fragment recovery of the single-stranded protocol is length-biased, with increased conversion of short fragments observed relative to double-stranded libraries. Although the efficiency of single-stranded DNA ligation may decrease for longer molecules (Li & Weeks 2006), we do not consistently observe such an effect at the 75bp maximum read length considered here, although this factor may become important for samples containing longer fragments (Gansauge & Meyer 2013). Overall, our results suggest that for maximising library complexity, the Dabney DNA isolation protocol used in conjunction with the single-stranded library protocol is optimal. Since effects on final library complexity are multiplicative, these protocols in combination could result in a more than 300-fold increase in DNA recovery compared to either of the other two DNA isolation methods combined with double-stranded library preparation. This single factor is likely to be decisive for many studies on ancient samples utilising either direct shotgun sequencing or hybridisation capture as downstream applications.

The second key factor determining data production efficiency is the total amount of usable sequence data generated per unit of sequencing effort. Unsurprisingly, we found that protocols associated with the recovery of longer fragment lengths generated greater amounts of data. For example, considering only merged reads, the double-stranded library generated significantly more usable data than the single-stranded protocol, and considering both merged and unmerged reads, the Rohland DNA isolation protocol in conjunction with the double-stranded library protocol provided the highest total data amounts. However, since these shifts to longer fragment lengths appear to result from decreased short fragment recovery, usable data production and complexity for a particular set of methods are directly opposed.

Researchers must therefore prioritise one of these two parameters before any assessment of protocol suitability can be made.

A final factor determining the efficiency of data production is that of endogenous content, which is highly variable among samples (Hofreiter *et al.* 2014). If the length distribution of endogenous and contaminant exogenous DNA are disparate, it is feasible that biases towards particular fragment lengths could result in a relative increase (or decrease) in endogenous DNA, which likely explains previous reports of changes in endogenous content associated with particular DNA isolation (Gamba *et al.* 2016) and library preparation (Bennett *et al.*2014; Wales *et al.* 2015) protocols. However, reliably predicting such effects is challenging without prior knowledge of relative sizes of target and non-target fractions within a sample.

## CONCLUSIONS

It makes intuitive sense that methods employed in the laboratory will have effects on high-throughput sequencing data. However, the characterisation of these effects within the context of palaeogenomic studies has received insufficient attention. Our study shows that two essential stages of sample preparation – DNA isolation and library preparation – can substantially influence total DNA recovery, introduce fragment length biases, and alter nucleotide composition. Moreover, we find that through the course of sample preparation, protocols used in these successive stages can have significant interaction effects. Such effects may reduce the predictive power of investigations of laboratory protocols in future experiments where other aspects of sample preparation are changed. Overall, careful consideration of the influence of laboratory methods is critical for any palaeogenomic study, both during the design of laboratory experiments and during analysis of the resulting data.

This latter is of particular relevance to studies involving mixed datasets derived using different methodologies.

Selecting the best laboratory methods is sample-and question-dependent, which precludes any generalised recommendations. Researchers are therefore advised to consider the properties of the sample and the relative importance of different data qualities when selecting appropriate methods. An accurate predictive model of expected results from alternative methodologies given specific sample properties is not possible based on the current data, but would represent an ideal solution.

## ACKNOWLEDGEMENTS

We thank Guy Bar-Oz for fieldwork and provision of samples, and Daniel Förster for useful discussion on data analysis. This work was supported by ERC consolidator grant No. 310763 GeneFlow to M.H.

## DATA ACCESSIBILITY

Sequence Datasets 1-4 used for statistical comparisons have been uploaded to Dryad.

## AUTHOR CONTRIBUTIONS

AB, GF, LD and MH designed and conceived of the study; AB and GF performed labwork; AB analysed data; AB, JP and MH interpreted results and wrote the manuscript. RP, BG, GR and CF provided samples. All authors gave final approval for publication.

